# Construction of invariant features for time-domain EEG/MEG signals using Grassmann manifolds

**DOI:** 10.1101/2024.03.11.584366

**Authors:** Rikkert Hindriks, Thomas O. Rot, Michel J.A.M. van Putten, Prejaas Tewarie

## Abstract

A challenge in interpreting features derived from source-space electroencephalography (EEG) and magnetoencephalography (MEG) signals is residual mixing of the true source signals. A common approach is to use features that are invariant under linear and instantaneous mixing. In the context of this approach, it is of interest to know which invariant features can be constructed from a given set of source-projected EEG/MEG signals. We address this question by exploiting the fact that invariant features can be viewed as functions on the Grassmann manifold. By embedding the Grassmann manifold in a vector space, coordinates are obtained that serve as building blocks for invariant features, in the sense that all invariant features can be constructed from them. We illustrate this approach by constructing several new bivariate, higher-order, and multidimensional functional connectivity measures for static and time-resolved analysis of time-domain EEG/MEG signals. Lastly, we apply such an invariant feature derived from the Grassmann manifold to EEG data from comatose survivors of cardiac arrest and show its superior sensitivity to identify changes in functional connectivity.

**Author Summary:** Electroencephalography (EEG) and magnetoencephalography (MEG) are techniques to non-invasively measure brain activity in human subjects. This works by measuring the electric potentials on the scalp (EEG) or the magnetic fluxes surrounding the head (MEG) that are induced by currents flowing in the brains’ grey matter (the ”brain activity”). However, reconstruction of brain activity from EEG/MEG sensor signals is an ill-posed inverse problem and, consequently, the reconstructed brain signals are linear superpositions of the true brain signals. This fact complicates the interpretation of the reconstructed brain activity. A common approach is to only use features of the reconstructed activity that are invariant under linear superpositions. In this study we show that all invariant features of reconstructed brain signals can be obtained by taking combinations of a finite set of fundamental features. The fundamental features are parametrized by a high-dimensional space known as the Grassmann manifold, which has a rich geometric structure that can be exploited to construct new invariant features. Our study advances the systematic study of invariant properties of EEG/MEG data and can be used as a framework to systematize and interrelate existing results. We use the theory to construct a new invariant connectivity measure and apply it to EEG data from comatose survivors of cardiac arrest. We find that this measure enables superior identification of affected brain regions.

## 1 Introduction

Electroencephalographic (EEG) and magnetoencephalographic (MEG) signals arise from current sources within the brain’s gray matter and are related to these sources by Maxwell’s equations [1]. Because Maxwell’s equations are linear, EEG/MEG sensor signals are (nearly) instantaneous linear mixtures of the underlying source signals and this fact complicates their interpretation of terms of the distribution of underlying current sources [2]. This particularly applies to *functional connectivity*, which refers to statistical dependencies between two or more brain signals [3] and is one of the fundamental concepts in systems neuroscience. Thus an observed statistical dependence between the signals measured at two different EEG/MEG sensors generally does not imply a statistical dependence between the signals from the brain regions underneath the sensors [4]. In other words, functional connectivity analysis of sensor EEG/MEG data is susceptible to spurious functional connectivity. Invasively recorded electrophysiological signals, such as cortical surface and local field potential recordings, are subject to the same mixing problem, although to a much lesser extent [5, 6, 7, 8, 9].

This mixing problem is usually dealt with in two steps. First, the EEG/MEG sensor signals are used to reconstruct the underlying source signals in the grey matter by inverting an EEG/MEG forward model. This is referred to as *inverse modeling* or *source reconstruction* [1, 10, 11]. Because the EEG/MEG inverse problem is ill-posed [1], inverse methods typically do not completely unmix the source signals and so the reconstructed source signals are still instantaneous linear mixtures of the true source signals, although the mixing is less severe. Residual mixing of source signals is occasionally referred to as *signal leakage* [2] and complicates the assessment of functional connectivity. A common second step, therefore, is to analyze the reconstructed source signals by using connectivity measures that are invariant under instantaneous linear mixing. Several such measures have been proposed, including the imaginary coherence [12], the lagged coherence [13], the (weighted) phase-lag index [14, 15], symmetrized phase-modulation functions [16], and the multivariate interaction measure [17] to name just a few.

Within the context of this two-step approach, a relevant question is whether there are more invariant connectivity measures and if there is a systematic way of obtaining them all. This question has been addressed only for stationary Gaussian frequency-domain signals [17]. In the current study, we address this question for arbitrary time-domain signals, thus including non-stationary and non-Gaussian signals. In other words, we construct a finite set of invariant connectivity measures from which *all* other invariant connectivity measures can be constructed by appropriately combining them. Although we are mainly interested in measures of functional connectivity, there exist invariant measures that characterize other aspects of multivariate signals and that have not yet been used to analyze EEG/MEG signals. Examples of such measures are the multivariate skewness and kurtosis measures proposed in [18] which relate to multivariate non-Gaussianity. Instead of the term *measure* we will use the more broadly used term *feature*. Thus, a feature is any real-valued function of a given EEG/MEG data-set.

The first goal of our study is to obtain a finite set of invariant features in terms of which all other invariant features can be constructed. In mathematical invariant theory, such features are referred to as *fundamental*. Our second goal is to use these fundamental features to construct new invariant connectivity measures. Besides bivariate connectivity measures, we construct higher-order and multidimensional connectivity measures. The term *higher-order connectivity* refers to statistical dependencies between more than two signals that cannot be reduced to pairwise dependencies. So *k*-th order connectivity refers to the connectivity between *k* signals. Third- and fourth-order connectivity have been observed in spike-train data [19, 20] and have recently received increased attention due to the discovery of discontinuous phase transitions in complex systems with higher-order interactions [21]. As far as we are aware, higher-order brain connectivity has not yet been investigated with EEG/MEG or extracellular electrophysiology. The term *multidimensional connectivity* refers to functional connectivity between multivariate signals [17, 22, 23] and can be used, for instance, to assess connectivity between regions-of-interest, each comprising *k* signals. We will construct both dynamic and static versions of such measures.

Let *X* be an *n* × *k* EEG/MEG data matrix of rank *k*, where *n* is the number of time points and *k* is the number of observed (i.e. reconstructed) source signals involved in the analysis. The construction of a fundamental set of invariant features for *X* is based on the following basic observation. The columns of *X* can be viewed as vectors in *n*-dimensional Euclidean space ℝ^*n*^. Due to residual mixing, each of these vectors is a linear combination of the true, but unknown, source signals. The observed and true signals, however, span the same *k*-dimensional subspace of ℝ^*n*^. Both the observed and true signals can therefore be viewed as bases for this subspace and the effect of mixing is a change of basis. In other words, invariant features of *X* are those that are independent of the chosen basis and thus only depend on the subspace *itself*, that is, on how the subspace is situated in ℝ^*n*^. It is therefore natural to construct features in EEG/MEG data matrices directly in terms of subspaces as this ensures invariance under mixing. This observation reduces the problem of finding all invariant features of *X* to finding all functions on the space of *k*-dimensional subspaces of ℝ^*n*^. This space is known as the *Grassmann manifold* [24].

In Section 2 we illustrate the general approach by using a simple example, provide the necessary mathematical background, and introduce the Grassmann manifold. In section 3 we discuss the Grass-mann approach to the EEG/MEG mixing problem and give a formal definition of invariant features. In Section 4 we describe two fundamental sets of invariant features. In Section 5 we use the fundamental features to construct new bivariate, higher-order, and multidimensional features, and features based on spatial activation patterns. As a proof of concept, we further apply one of our invariant features to EEG data from comatose survivors of cardiac arrest (Section 6), and demonstrate that these invariant features are more sensitive to disease-induced effects than conventional functional connectivity metrics.

## 2 Preliminaries

### 2.1 Illustrative example

In this section, we consider the simplest scenario in which we can define a non-trivial property of EEG data that is invariant under instantaneous linear mixing. Although simple, it contains most of the mathematical concepts that are needed to treat the general scenario. In Section 2.2 we will introduce these concepts in a more formal way.

Let *x* = (*x*_1_, *x*_2_)^T^ ∈ ℝ^2^ be a reconstructed time-domain EEG signal from a given location within the brain and assume that either *x*_1_ or *x*_2_ is non-zero. Note that the signal is very short; it only comprises two time points. Due to signal leakage, we only know *x* up to an unknown non-zero scalar *a*. In other words, if 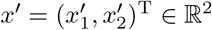 is the true (but unknown) signal, then *x* = *ax*′ with *a* unknown. Is there anything that can be said about the true signal on the basis of the reconstructed signal? Let *f* (*x*) be a feature of the observed signal. A feature contains information about the true signal only if *f* (*ax*) = *f* (*x*) for all signals *x* and non-zero scalars *a*. We refer to such a feature as *invariant* under mixing. An example of a feature that is *not* invariant is the power of 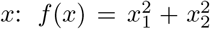, because *f* (*ax*) = *a*^2^*f* (*x*) ≠ *f* (*x*). An example of a feature that *is* invariant, is the power ratio 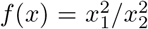, because 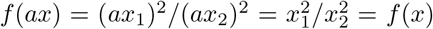. Note that the power ratio can only be calculated if *x*_2_ ≠0.

Because of mixing, we do not know *x* exactly, only that it lies on the line through *x* and the origin in ℝ^2^. We denote this line by [*x*]. Because we do not know where *x* lies on [*x*], no information is lost when working with [*x*] instead of with *x*. So there is a correspondence between the set of signals that cannot be distinguished from each other due to mixing and lines in ℝ^2^. In other words, by working with lines instead of with signals, the redundancy in the presentation of the signals that is introduced by signal leakage is removed. We denote the space of lines through the origin in ℝ^2^ by ℝ*P* ^1^. Invariant features correspond to functions *f* : ℝ*P* ^1^ → ℝ. How can we exploit this observation to obtain all invariant features of *x*?

The key property of ℝ*P* ^1^ is that it is one-dimensional, in the sense that it can locally be parametrized by a single coordinate. Such a set is called a *one-dimensional manifold*. The one-dimensional manifold ℝ*P* ^1^ is known as the *projective line*. For example, the non-horizontal line that is determined by the point (*X, Y*)^T^ ∈ ℝ^2^ with *Y*≠ 0 can be mapped to ℝ by the function *ϕ*(*X, Y*) = (*X/Y*, 1) and this establishes a correspondence between the subset of ℝ*P* ^1^ consisting of non-horizontal lines and (a copy of) ℝ. The coordinate of the line hence is *X/Y*. Figure 1 provides an illustration. This implies that any function on ℝ*P* ^1^ can be written as a function of the single coordinate *X/Y*, at least locally. In terms of the EEG signal *x* = (*x*_1_, *x*_2_)^T^ it means that for *x*_2_ ≠ 0, any invariant feature is a function of *x*_1_*/x*_2_. Examples of invariant features hence are *f* (*x*_1_, *x*_1_) = (*x*_1_*/x*_2_)^2^ and *f* (*x*_1_, *x*_2_) = atan(*x*_2_*/x*_1_). However, essentially there is only one independent invariant feature, namely the ratio *x*_1_*/x*_2_.

**Figure 1.**
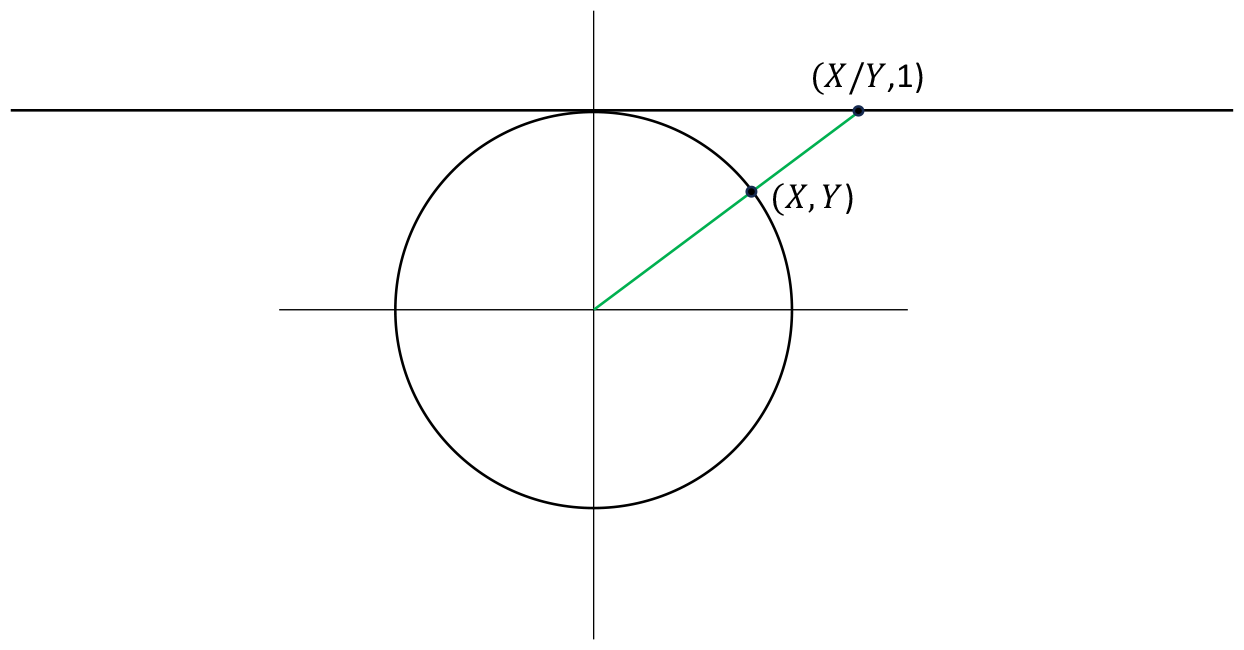
Local coordinates on the projective line. Shown in green is the non-horizontal line in ℝ^2^ (i.e. a “point” on the projective line) that is determined by the point (*X, Y*) ∈ ℝ^2^ on the unit circle (i.e. *X*^2^ + *Y* ^2^ = 1) and with *Y*≠ 0, together with its image (*X/Y*, 1) under the function *ϕ*, which is located on the horizontal line at height 1. The function *ϕ* establishes a correspondence between non-horizontal lines in ℝ^2^ and points on the horizontal line at height 1. The number *X/Y* is the coordinate of the green line. The coordinates are local because they are not defined for all lines in ℝ^2^, but only for non-horizontal lines.

A drawback of the above approach to finding all invariant features of the signal *x* is that any parametrization of the projective line is necessarily local, because the projective line is topologically distinct from the real line. Another drawback is that any point on the projective line has many different parametrizations and making a choice requires singling out a particular time point of *x* (in the above case the second time-point *x*_2_) which is unnatural in most experimental contexts. In this study we therefore focus on a different approach, which is to embed the projective line in a vector space *V* and use the vector space coordinates to obtain global coordinates. These global coordinates will then serve as building blocks for invariant features as explained below.

Which vector space *V* and embedding *p* : ℝ*P* ^1^ → *V* should we use? We first note that lines through the origin in ℝ^2^ are one-dimensional linear subspaces of ℝ^2^ and that these correspond to 2 × 2 projection matrices of rank one: With every line in ℝ^2^ we can associate a unique projection matrix of rank one, namely the matrix of the orthogonal projection onto that line. Also, with every such matrix we can associate a unique line, namely the line onto which the matrix projects. So we take *V* to be the vector space of symmetric 2 × 2 matrices and let *p*(*x*) be the orthogonal projection matrix onto the line [*x*]:

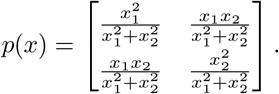

This embedding allows to identify the projective line with its image under the function *p*, which is the (non-linear) subspace of *V* consisting of projection matrices of rank one. Note that because the matrix *p*(*x*) is symmetric, the vector space *V* is three-dimensional, and hence we obtain the following coordinate function *c* : ℝ*P* ^2^ → ℝ^3^:

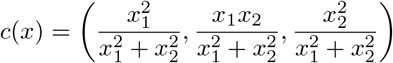

Note that the coordinates of *c*(*x*) are indeed invariant under scaling of *x* by non-zero numbers *a*. Furthermore, every invariant feature of *x* can be build from these three coordinates and every function of the coordinates of *c*(*x*) is an invariant feature of *x*. For example, the invariant feature 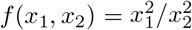 can be build by dividing the first coordinate of *c*(*x*) by its third coordinate. We do note that the coordinates of *c*(*x*) are not independent. Indeed, since the projective plane is one-dimensional, there can only exist one independent invariant, whereas *c*(*x*) has three coordinates. So there must be two independent relations between the coordinates of *c*(*x*). Indeed, we see that *c*_1_(*x*)*c*_3_(*x*) = *c*_2_(*x*)^2^ and *c*_1_(*x*) + *c*_2_(*x*) = 1.

### 2.2 Mathematical background

In this section we list several mathematical concepts that are used in later sections. Each concept is illustrated by the example from Section 2.1 and some further examples that are relevant for the study of invariant features of EEG/MEG data matrices.

#### EEG/MEG data frame

An *EEG/MEG data frame* (or simply a *frame*) is an *n* × *k* matrix whose columns are linearly independent EEG/MEG signals of length *n* from *k* different locations within the brain (so *k* ≤ *n*). A frame is denoted by *X*. We also refer to *X* as a *k*-frame in ℝ^*n*^. A frame is called *orthonormal* if its columns have unit norm and are orthogonal or equivalently if *X*^T^*X* = *I*_*k*_, where *I*_*k*_ is the *k* ×*k* identity matrix. Figure 2 provides an illustration of a general and an orthonormal frame.

**Figure 2.**
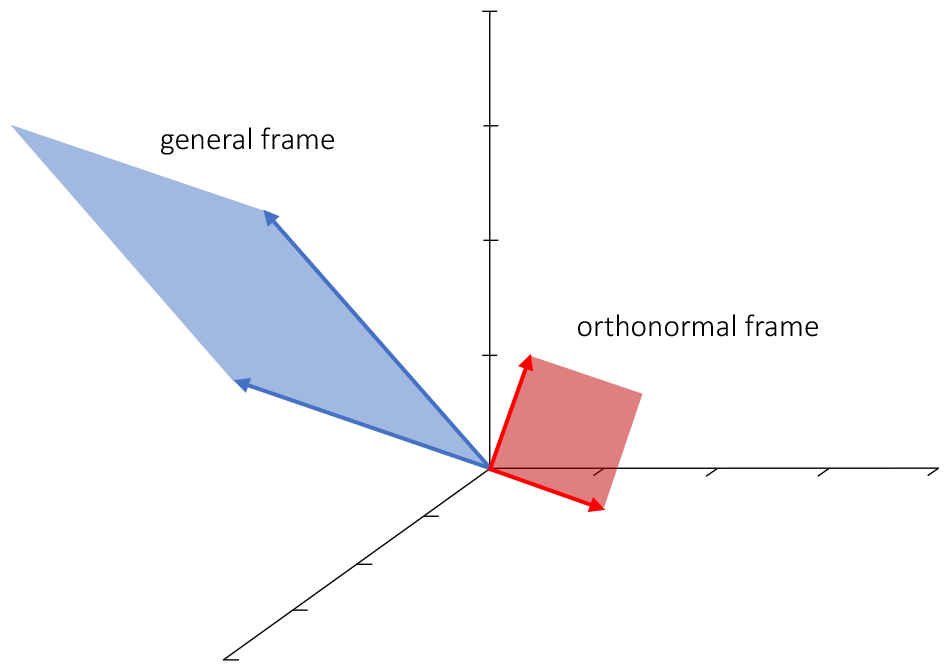
EEG/MEG data frames. Shown are two 2-frames (red and blue vector-pairs) in 3-dimensional Euclidean space ℝ^3^. The vectors constituting each frame correspond to EEG/MEG signals of length 3. The blue frame is a general frame and the red frame is an orthonormal frame. The planes spanned by the frames are illustrated by the red and blue rectangles.

*Example:* For *n* = 2 and *k* = 1 we obtain the example from Section 2.1. More generally, for *k* = 1, frames correspond to EEG/MEG signals of length *n*.

#### Canonical angles and correlations

Let *V*_1_ and *V*_2_ be *k*_1_- and *k*_2_-dimensional subspaces of ℝ^*n*^ and let *m* = min{*k*_1_, *k*_2_}. The *canonical angles θ*_1_, …, *θ*_*m*_ between *V*_1_ and *V*_2_ completely characterize the relative orientations of *V*_1_ and *V*_2_ in ℝ^*n*^ and are defined recursively as follows. The first canonical angle *θ*_1_ is defined by

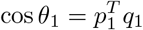

where *p*_1_ ∈ *V*_1_ and *q*_1_ ∈ *V*_2_ are unit vectors that maximize the inner product 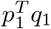. The second principal angle *θ*_2_ is defined by

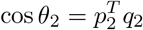

where *p*_2_ ∈ *V*_1_ and *q*_2_ ∈ *V*_2_ are unit vectors that maximize the inner product 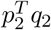 and are orthogonal to the first principal vectors (i.e. 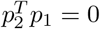 and 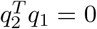). Continuing in this way we obtain *m* unique principal angles. Figure 3 provides an illustration.

**Figure 3.**
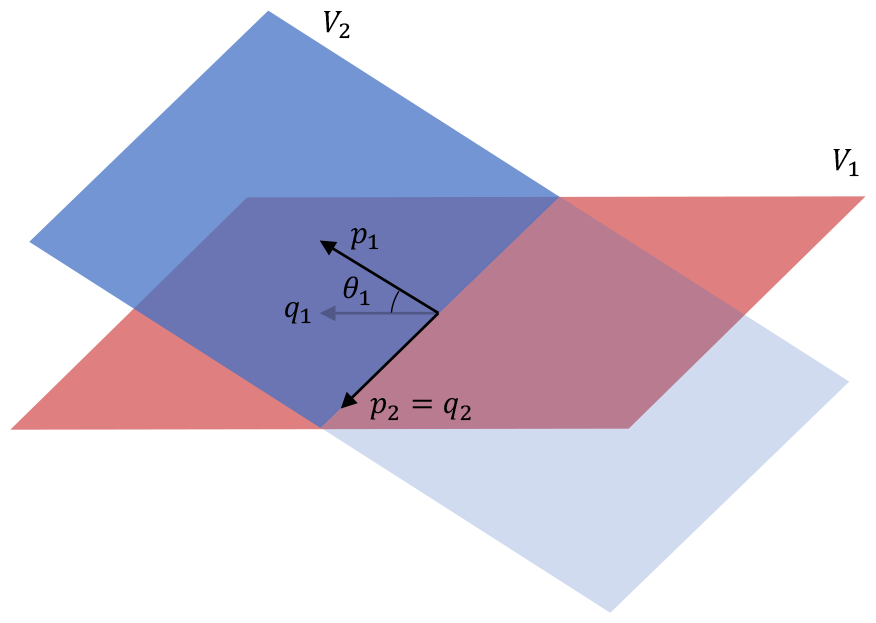
Canonical angles. Shown are two 2-dimensional subspaces *V*_1_ and *V*_2_ of ℝ^3^ (red and blue rectangles) together with their canonical angles *θ*_1_ and *θ*_2_. The second canonical angle is the angle between the vectors *p*_2_ and *q*_2_ and equals zero, because *p*_2_ = *q*_2_. This reflects the fact that *V*_1_ and *V*_2_ intersect in a line.

The canonical angles can be calculated by using the singular value decomposition as follows [25]. Take orthonormal frames *Y*_1_ and *Y*_2_ that span *V*_1_ and *V*_2_, respectively, and compute the singular value decomposition of the matrix 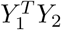:

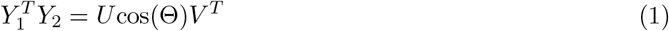

where *U* and *V* are, respectively *k*_1_ × *m* and *k*_2_ × *m* matrices with orthonormal columns, and cos(Θ) is a *m*-dimensional diagonal matrix that contains the singular values of 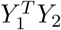. The notation cos(Θ) conveys that the singular values of 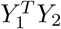 are the cosines of the canonical angles:

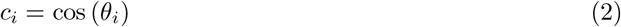

where *c*_*i*_ is the *i*-th singular value of 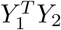.

Let *X*_1_ and *X*_2_ be *k*_1_- and *k*_2_-frames in ℝ^*n*^. The *canonical correlations* between *X*_1_ and *X*_2_ are the cosines of the canonical angles between the columns spans [*X*_1_] and [*X*_2_] of *X*_1_ and *X*_2_. They characterize the properties of the cross-covariance matrix 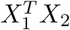 that are independent of the bases in which the frames are expressed. They can be calculated by whitening *X*_1_ and *X*_2_ to obtain orthonormal frames *Y*_1_ and *Y*_2_ (see below) and subsequently calculating the singular values of the cross-covariance matrix 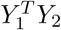 between the whitened frames.

*Example:* The canonical angle between two vectors *x*_1_ and *x*_2_ in ℝ^*n*^ coincides with the ordinary notion of angle and their canonical correlation coincides with the Pearson correlation coefficient.

#### Manifold

A *manifold* ℳ of dimension *n* is a space that locally looks like *n*-dimensional Euclidean space ℝ^*n*^. This means that for every point *p* ∈ ℳ there exists a neighbourhood *U* ⊆ ℳ and a map *ϕ* : *U* → *V* ⊆ ℝ^*n*^ to a subset *V* of ℝ^*n*^ that is one-to-one and onto. The image *ϕ*(*q*) of a point *q* ∈ *U* contains the local coordinates of *q*. Intuitively, one can think of *n*-dimensional manifolds as curved *n*-dimensional spaces. Figure 4 provides an illustration.

**Figure 4.**
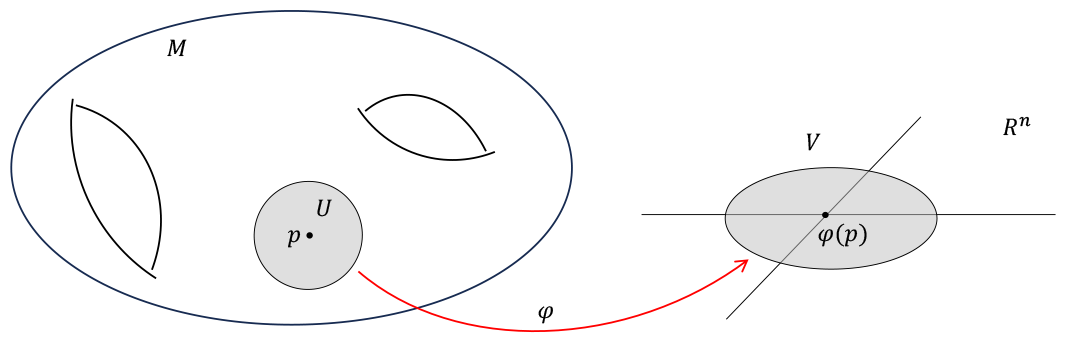
Manifold and local coordinates. Schematic illustration of an *n*-dimensional manifold ℳ, together with a point *p* ∈ ℳ and a surrounding neighborhood *U* ⊆ ℳ. The function *ϕ* (red arrow) maps *U* to a subset *V* of *n*-dimensional Euclidean space ℝ^*n*^. The inverse function *ϕ*^−1^ : *V* → *U* is a local parametrization of ℳ. Every point on the manifold has such a local parametrization and they endow the manifold with local coordinates.

*Examples:* Euclidean space ℝ^*n*^ is a manifold of dimension *n*. Other examples are the 1-dimensional sphere *S*^1^ from Section 2.1, which is a manifold of dimension 1, and the two-sphere *S*^2^, which is a manifold of dimension 2. The space of *k*-frames in ℝ^*n*^ is an *nk*-dimensional manifold called the *non-compact Stiefel manifold*. We will denote it by *F*_*n,k*_. The space of orthonormal *k*-frames in ℝ^*n*^ is a manifold of dimension *nk* − *k*(*k* + 1)*/*2 called the *compact Stiefel manifold*. We will denote it by 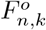. Note that 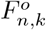 is contained in *F*_*n,k*_. The space of *k*-dimensional subspaces in ℝ^*n*^ is a compact manifold of dimension *k*(*n* − *k*) called the *Grassmann manifold*. We will denote it by *G*_*n,k*_. The Stiefel manifold can be mapped to the Grassmann manifold by the map *π* : *F*_*n,k*_ → *G*_*n,k*_ defined by *π*(*X*) = [*X*], where [*X*] denotes the span of the columns of *X*. The pre-image *π*^−1^(*V*) of a *k*-plane *V* consists of all *k*-frames that span *V*.

#### Whitening transformation

A *whitening transformation* is a map *ϕ* : 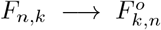from the non-compact Stiefel manifold to the compact Stiefel manifold of the form *ϕ*(*X*) = *XW* for some invertible *k* × *k* matrix *W* and which restricts to the identity map on 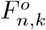. The transformation *W* will depend on *X*, but we suppress this from the notation. In the context of EEG/MEG signals, whitening decorrelates the signals in *X* and standardizes their variances to 1. A whitening transformation always exists and can be constructed, for instance, by taking a square root of the spatial covariance matrix *X*^*T*^ *X* of *X*.

#### Group

A *group* is a set *G* together with a product operation. The product operation *G* × *G* → *G* maps a pair (*a, b*) of group elements to another group element *ab* called the *product* of *a* and *b*. The product is assumed to have the following three properties: (i) (*ab*)*c* = *a*(*bc*), (ii) there is an element *e* ∈ *G* such that *ea* = *ae* = *e* for all *a* ∈ *G* (identity element), and (iii) for every *a* ∈ *G* there is a *a*′ ∈ *G* such that *aa*′ = *a*′*a* = *e*. The element *a*′ is called the *inverse* of *a* and is denoted by *a*^−1^.

*Examples:* The group R*/*{0} consists of non-zero real numbers with ordinary multiplication (*a, b*) ↦ *ab* (see Section 2.1). The identity is 1. Another example is the group of invertible *k* × *k* matrices with real entries, together with matrix multiplication (*A, B*) ↦ *AB*. This group is called the *general linear group* and is denoted by GL_*k*_. Yet another example is the set of *k* × *k* orthogonal matrices with real entries, again with matrix multiplication. It is called the *orthogonal group* and we denote it by O_*k*_.

#### Group action

Let *G* be a group and let 𝒳 be an arbitrary set. *G* is said to *act* on 𝒳 if every *g* ∈ *G* induces a bijection *L*_*g*_ : 𝒳 → 𝒳 from 𝒳 to itself, such that *L*_*e*_ is the identity map on 𝒳 and *L*_*h*_ ° *L*_*g*_ = *L*_*hg*_, where ° denotes concatenation of maps.

*Examples:* Let 𝒳 be the non-compact Stiefel manifold *F*_*n*,1_ of 1-frames in ℝ^*n*^ (i.e. the manifold of EEG signals of length *n*). The group ℝ*/*{0} acts on *F*_*n*,1_ by scalar multiplication: *L*_*g*_(*x*) = *ax*. This describes the effect of mixing in the example from Section 2.1. More generally, the general linear group GL_*k*_ acts on *F*_*n,k*_ by right-multiplication: *L*_*A*_*X* = *XA*. Similarly, the orthogonal group O_*k*_ acts on the compact Stiefel manifold 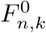 by right-multiplication.

#### Orbit

Let *G* be a group acting on a set 𝒳 and let *x* ∈ 𝒳 be an element of 𝒳. The subset of 𝒳 that is traced out by letting elements of *G* act on *x* is called the *orbit* of *x*. It is the set

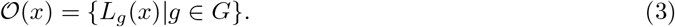

*Examples:* The orbit of an EEG signal *x* ∈ *F*_*n*,1_ under the above action of ℝ*/*{0} is the line in ℝ^*n*^ that is spanned by *x*, excluding the origin. More generally, the orbit of a *k*-frame *X* in ℝ^*n*^ under the action of the general linear group GL_*k*_ on the non-compact Stiefel manifold *F*_*n,k*_ is the *k*-plane [*X*] that is spanned by the columns of *X*, again excluding the origin.

#### Orbit space

Let *G* be a group acting on a space 𝒳. The space of all orbits of *G* is called the *orbit space* and is denoted by 𝒳 */G*. So points on 𝒳 */G* correspond to orbits. There is a map *π* : 𝒳 → 𝒳 */G* that maps *x* to its orbit [*x*].

*Examples:* The orbit space of the action of ℝ*/*{0} on *F*_2,1_ is the projective line R*P* ^1^ (see Section 2.1). More generally, the orbit space of ℝ*/*{0} acting on *F*_*n*,1_ is (*n* − 1)-dimensional projective space ℝ*P* ^*n*−1^. Even more generally, the orbit space of the action of GL_*k*_ on *F*_*n,k*_ is the Grassmann manifold *G*_*n,k*_. So points on *G*_*n,k*_ correspond to *k*-planes in ℝ^*n*^. The Grassmann manifold is also equal to the orbit space of the action of the orthogonal group O_*k*_ on the compact Stiefel manifold 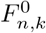:

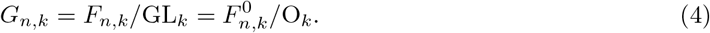

An explicit bijection between these two representations of the Grassmann manifold is the map that sends an orbit [*X*] in *F*_*n,k*_*/*GL_*k*_ to the orbit of a whitened version *Y* = *XW* of *X* in 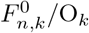.

#### Alternating tensor

Let *V* be an *n*-dimensional vector space. After choosing a bases, *V* can be identified with ℝ^*n*^. A *k*-tensor *T* on *V* can now be thought of as a *k*-dimensional array with sides of length *n*. The entries of *T* are the coordinates of the tensor with respect to the chosen basis. We denote the entries of *T* by 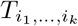 where each of the indices runs from 1 to *n*. So a 1-tensor on ℝ^*n*^ is an *n*-dimensional vector with entries 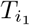 for *i*_1_ = 1, …, *n* and a 2-tensor on ℝ^*n*^ is an *n* × *n* matrix with entries 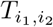 for *i*_1_, *i*_2_ = 1, …, *n*, etc. Any permutation *σ* of the set of *k* letters can be written as the concatenation of transpositions and the sign of *σ* is sgn(*σ*) = (−1)^*m*^, where *m* is the number of transpositions. A permutation is *odd* if its sign is negative and *even* if its sign is positive. A *k*-tensor *T* on ℝ^*n*^ is *alternating* if, under permutation of its subscripts, it changes sign according to the sign of the permutation:

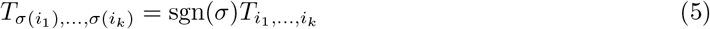

for all permutations *σ* of *k* letters. Alternating *k*-tensors on ℝ^*n*^ form a vector space of dimension 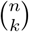 which is commonly denoted by ⋀^*k*^ ℝ^*n*^.

*Examples*. Alternating 2-tensors on ℝ^*n*^ satisfy *T*_*j,i*_ = −*T*_*i,j*_ and hence are the same as anti-symmetric *n*-dimensional matrices. Similarly, alternating 3-tensors on ℝ^*n*^ satisfy *T*_*jik*_ = *T*_*ikj*_ = *T*_*kji*_ = −*T*_*ijk*_ and *T*_*kij*_ = *T*_*jki*_ = *T*_*ijk*_.

#### Wedge product

Let *x*_1_, …, *x*_*k*_ be vectors in ℝ^*n*^. With these vectors we can associate an alternating *k*-tensor on ℝ^*n*^ which is denoted by *x*_1_ ∧ · · · ∧ *x*_*k*_. Its entries are given by

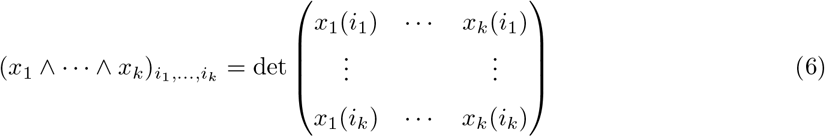

for *i*_1_, …, *i*_*k*_ ranging between 1 and *n* and where *x*_*j*_(*m*) is the *m*-th entry of *x*_*j*_. The magnitude of *x*_1_ ∧ · · · ∧ *x*_*k*_ (i.e. the square root of the sum of its squared entries) equals the *k*-dimensional volume of the parallelpiped spanned by *x*_1_, …, *x*_*n*_.

*Example:* Let *k* = 2 and *n* = 3. Let *x* = (*x*_1_, *x*_2_, *x*_3_)^T^ and *y* = (*y*_1_, *y*_2_, *y*_3_)^T^ be column vectors. Their wedge product is the following 2-tensor on ℝ^3^:

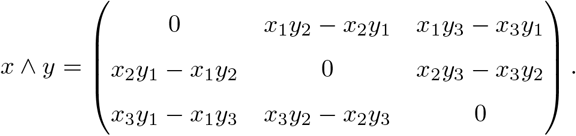

Note that *x* ∧ *y* is an anti-symmetric matrix and hence is determined by its three upper-triangular entries. These entries can be placed in a 3-dimensional vector and form the cross-product between *x* and *y*:

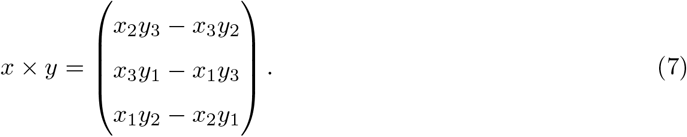

Thus, the wedge product is a generalization of the cross-product to more than two vectors of higher dimension. The magnitude of the cross-product *x* × *y* is the surface area (i.e. the 2-volume) of the parallelogram (i.e. a 2-dimensional parallelpiped) spanned by *x* and *y*.

#### Projectivation of a vector space

Let *V* be an *n*-dimensional real vector space. The *projectivation* of *V* is the space of lines through the origin in *V*. It is obtained by identifying each non-zero vector *v* ∈ *V* with its scalar multiples *av* for *a* ∈ ℝ*/*{0}. We will denote the projectivation of *V* by 𝒫 (*V*).

*Examples:* The projectivation of ℝ^3^ is the space of lines in ℝ^3^. Since lines in ℝ^3^ correspond to pairs of antipodal points on the unit sphere *S*^2^ as any antipodal pair uniquely determines a line through the origin, the projectivation of ℝ^3^ is the projective plane ℝ*P* ^2^, which also equals *G*_3,1_. Topologically, it is obtained by taking *S*^2^ and ”gluing” (i.e. identifying) antipodal points. Figure 5 provides an illustration.

**Figure 5.**
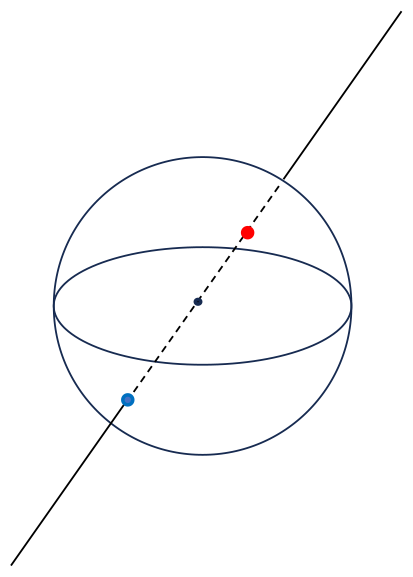
Projectivation of ℝ^3^. Shown is the unit 2-sphere *S*^2^, together with a line through the origin of ℝ^3^ and the intersection of the line with a pair of antipodal points on *S*^2^ (red and blue dots). Points on the projectivation P(ℝ^3^) of ℝ^3^ correspond to pairs of antipodal points.

## 3 A geometric approach to the EEG/MEG mixing problem

### 3.1 The EEG/MEG mixing problem

In this section we describe the mixing problem for EEG/MEG data matrices, give a geometric inter-pretation of it, and explain the relation to the Grassmann manifold.

Let *X*′ be a real *n* × *k* matrix whose columns carry the true source signals from *k* locations within the brain. Here *n* ≥ *k* is the number of time-points. We assume that the signals are linearly independent so that *X*′ has rank *k*. We refer to an *n* × *k* matrix of rank *k* as a *k-frame* in ℝ ^*n*^ (see Section 2.2). Let *X* be the real *n* × *k* matrix whose columns carry the observed (i.e. reconstructed) source signals from the same *k* brain locations. Due to residual mixing, the observed signals are instantaneous linear mixtures of the true signals:

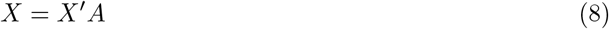

where *A* is a *k* × *k* matrix with real entries, which we will refer to as the *mixing matrix*. The mixing problem for source-space EEG/MEG data matrices is that we do not observe *X*′, but only *X*.

Two main approaches to the mixing problem are the following. The first is recent and is restricted to situations in which the mixing matrix is known. This is the case if the projection from sensor-space to source-space is carried out by a linear transformation such as the minimum norm solution, low-resolution electrical tomography, or a linearly constrained minimum variance beamformer [26, 27, 10]. The mixing matrix then coincides with the resolution matrix associated with the inverse transformation [28, 29] and this fact can be used to define invariant connectivity measures that depend on the mixing matrix [30, 31, 29, 32]. In this approach, the mixing matrix is not required to be invertible. We will not discuss this approach in the current study.

The second approach does not require knowledge of the mixing matrix. The strategy is to define connectivity measures that are invariant under mixing by *all* possible mixing matrices. Examples of such measures where mentioned in Section 1 and include the imaginary coherence [12], the lagged coherence [13], the (weighted) phase-lag index [14, 15], symmetrized phase-modulation functions [16], and the multivariate interaction measure [17]. This approach requires the mixing matrix to be invertible, which can be a valid assumption, depending on which brain locations are included in the analysis. We focus on this second approach and our goal is to determine all possible invariant features of EEG/MEG data matrices, including functional connectivity measures. In the remainder of the text, we therefore assume that the mixing matrix *A* is invertible.

To explain the role of the Grassmann manifold in solving the EEG/MEG mixing problem, we first describe the mixing problem in geometric terms. The key observation is that, because the mixing matrix *A* is invertible, the columns of the observed frame *X* and those of the true frame *X*′ span the same *k*-dimensional subspace of ℝ ^*n*^. This is illustrated in Figure 6. A *k*-frame in ℝ ^*n*^ corresponds to a basis of a *k*-dimensional subspace (i.e. a *k*-plane) of ℝ ^*n*^ and the effect of mixing is to change to a different basis. Hence the properties of a *k*-frame that are invariant under mixing, are those that are independent of the choice of basis and thus only depend on how the *k*-plane is embedded in ℝ ^*n*^. Examples of invariant properties are the angles that the *k*-plane makes with the standard coordinates axes of ℝ ^*n*^.

**Figure 6.**
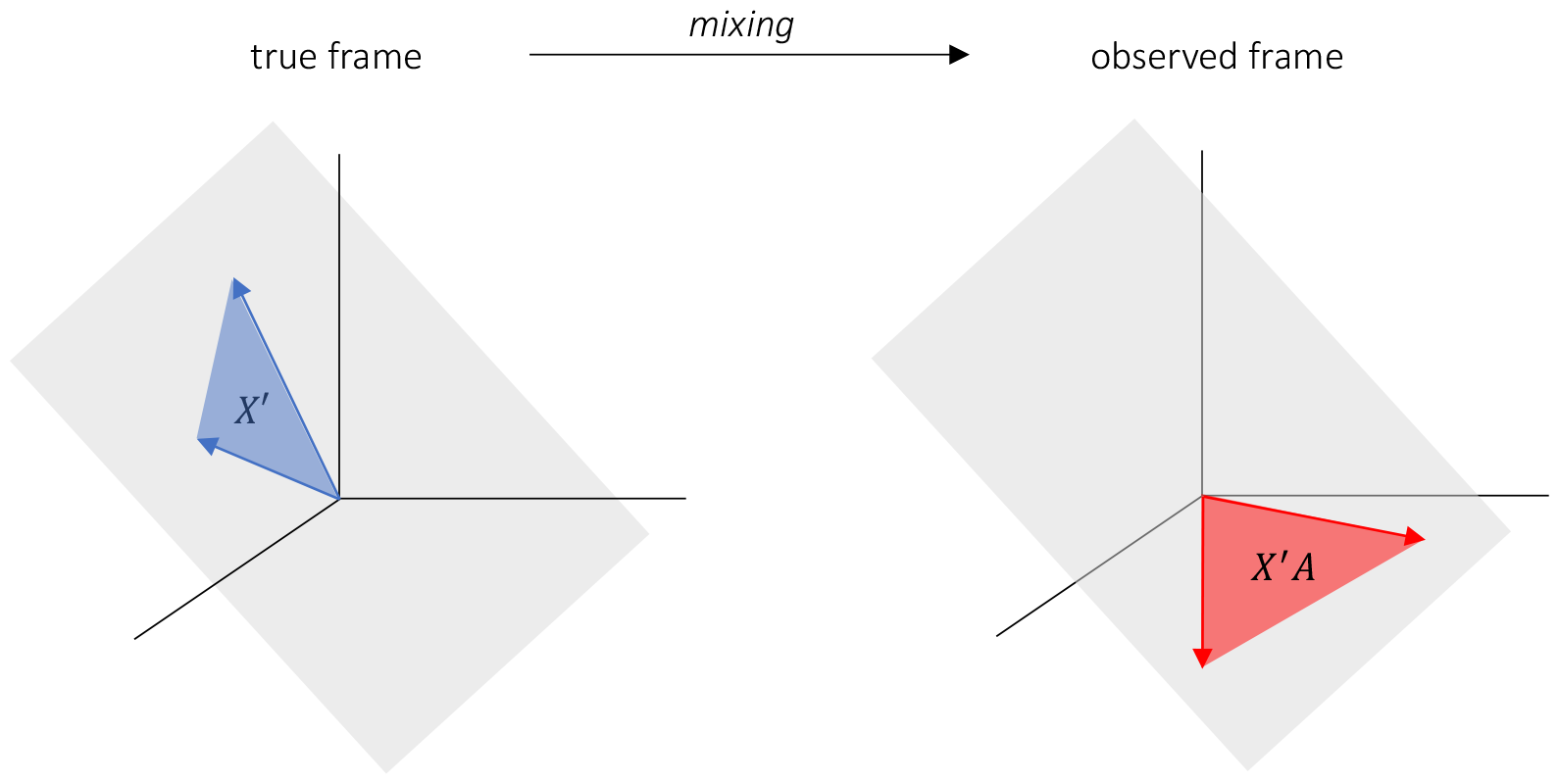
Effect of mixing on EEG data frames. Shown are a true frame *X*′ (blue) and the observed frame *X* = *X*′*A* (red) in *n*-dimensional Euclidean space ℝ^*n*^. The blue and red vectors correspond, respectively, to the true and observed source signals. The true and observed frames span the same *k*-dimensional subspace (grey plane through the origin) of ℝ^*n*^.

We now make the connection with the Grassmann manifold *G*_*n,k*_ of *k*-planes in ℝ ^*n*^. We first note that *k*-dimensional mixing matrices form the general linear group GL_*k*_ (see Section 2.2). Recall from Section 2.2 that EEG/MEG data frames correspond to points on the non-compact Stiefel manifold *F*_*n,k*_. With every mixing matrix *A* ∈ GL_*k*_ we can associate an invertible map *L*_*A*_ : *F*_*n,k*_ → *F*_*n,k*_ from the non-compact Stiefel manifold to itself, namely the map

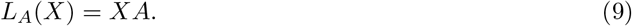

This map thus describes the effect of mixing on EEG/MEG data frames. It defines an action of the group GL_*k*_ on *F*_*n,k*_ (see Section 2.2). Also recall from Section 2.2 that the orbit 𝒪 (*X*) of an EEG/MEG data frame *X* ∈ *F*_*n,k*_ is the subspace of *F*_*n,k*_ that is traced out when letting GL_*k*_ act on *X*:

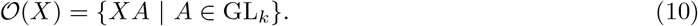

The orbit of *X* consists of those frames that have the same column space as *X*. In other words, the orbit space of *X* is the *k*-plane in ℝ ^*n*^ that is spanned by *X*. This plane is denoted by [*X*]. Because of mixing, EEG/MEG data frames that are in the same orbit cannot be distinguished and it is therefore more natural to work in the orbit space, rather than with frames. As explained in Section 2.2, the orbit space is the Grassmann manifold *G*_*n,k*_.

### 3.2 Mixing-invariant features

Informally, a feature of a *n* × *k* EEG data frame *X* ∈ *F*_*n,k*_ is any property of *X* that can be expressed by a single real number. Recall that the columns of *X* are the observed EEG signals from *k* different brain regions. Examples of features of individual signals are amplitude, power, peak frequency, entropy, and LZ complexity, among many others. Examples of features that depend on multiple signals of *X* are the Kuramoto order parameter, multivariate information measures, and all functional connectivity measures. Formally, a *feature* of an *n* × *k* EEG data frame *X* ∈ *F*_*n,k*_ is the image *f* (*X*) of a real-valued function *f* : *F*_*n,k*_ → ℝ. Note that in this definition we have assumed that a feature is defined for every data frame *X* ∈ *F*_*n,k*_. In this study, we are interested in features that are invariant under linear and instantaneous mixing and particularly in invariant measures of functional connectivity.

We refer to a feature *f* as *invariant* under linear and instantaneous mixing if

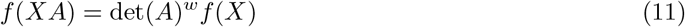

for all *A* ∈ GL(*k*) and *X* ∈ *F*_*n,k*_ and for some integer *w* called the *weight* of *f*. Following standard practice in invariant theory [33] we refer to features with zero and non-zero weights as *absolute* and *relative*, respectively. EEG studies mostly focus on absolute invariant features such as the imaginary part of coherence, the phase lag index, the (weighted) phase-lag index, symmetrized phase-modulation functions, the multivariate interaction measure, and the standardized imaginary covariance.

There are, however, several reasons for preferring the more general definition in Eq. (11). First, although absolute invariant features can be used as estimators for corresponding population quantities, relative invariant features can be used as statistics for testing statistical hypotheses. An example is the imaginary part of the cross-spectrum, which is an invariant feature with weight 2. Although it depends on the power of the signals and therefore is not a good measure of functional connectivity, it can be used in testing for significant imaginary coherence, because the imaginary coherence is zero if and only if the imaginary part of the cross-spectrum is zero. Second, from a mathematical perspective, including relative invariant features is more natural [33] and many absolute invariant features can be constructed by taking ratios of relative invariant features. An example is the lagged coherence, which is the ratio of two invariants of weight 2.

## 4 Fundamental invariant features

The Grassmann manifold *G*_*n,k*_ is a manifold of dimension *k*(*n* − *k*) and hence coordinates can be constructed in a neighborhood of every point, yielding *k*(*n* − *k*) independent invariant features. However, these features are only defined locally and therefore do not cover all EEG/MEG data frames. Furthermore, the standard way of calculating local coordinates requires selecting *k* arbitrary time-points, which is unnatural in most experimental settings. By embedding the Grassmann manifold in a vector space, the coordinates of the vector space can be restricted to the (embedded) Grassmann manifold and hence yield coordinates that are defined for all EEG/MEG data frames. This is illustrated in Figure 7.

**Figure 7.**
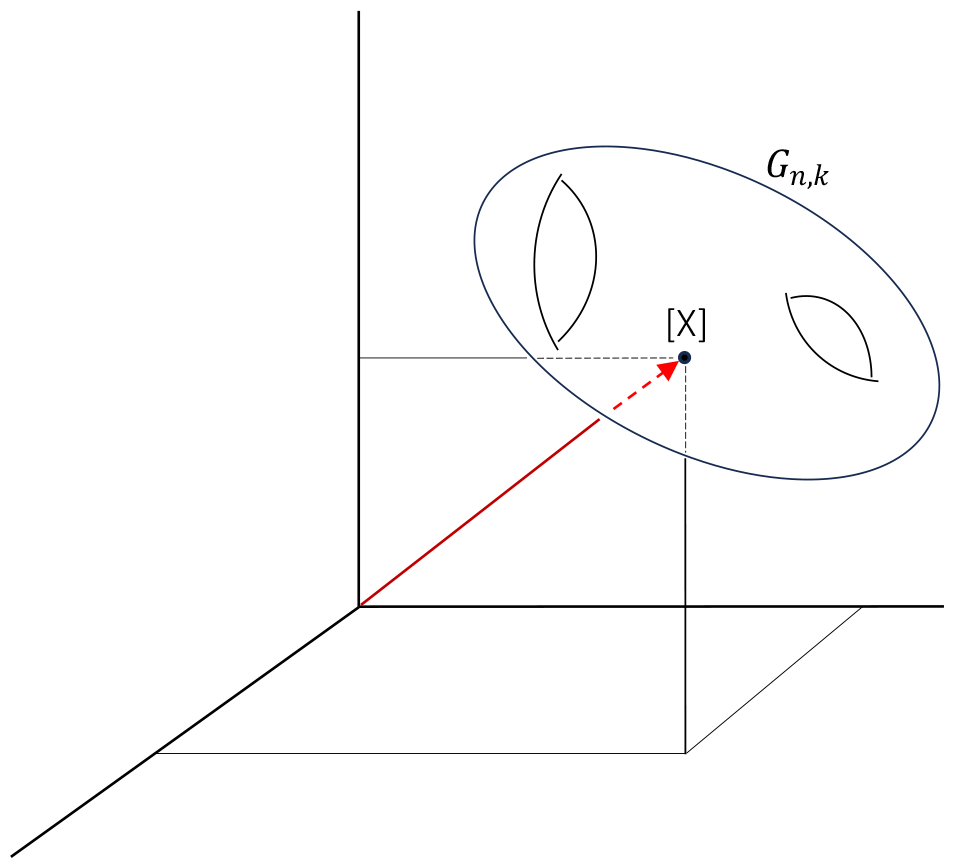
Embedding of the Grassmann manifold into a vector space. Shown is the Grassmann manifold *G*_*n,k*_ of *k*-planes in ℝ^*n*^ (curved surface) embedded into a vector space. Any point [*X*] on the Grassmann manifold (black dot) can be identified with a unique vector in the vector space (red arrow) and the coordinates of the vector can therefore be used as coordinates of [*X*]. In particular, any function on the Grassmann manifold (i.e. any invariant feature) can be expressed in terms of these coordinates. In this sense, these coordinates constitute building blocks for invariant features.

Recall from Section 3.2 that invariant features can be viewed as real-valued functions on the Grassmann manifold. Any invariant feature can thus be expressed in these coordinates and they therefore constitute building blocks for invariant features. In the context of EEG/MEG data, we will refer to these coordinates as *fundamental features*.

We consider two sets of fundamental features. The *fundamental affine features* are obtained by embedding the Grassmann manifold *G*_*n,k*_ in the vector space of symmetric *n* × *n* matrices, which gives *n*(*n* + 1)*/*2 fundamental features. The *fundamental projective features* are obtained by embedding the Grassmann manifold in (the projectivation of) the vector space of alternating *k*-tensors on ℝ ^*n*^, which gives 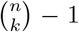 fundamental features. Because any point [*X*] on the Grassmann manifold can be represented by either affine or projective features, the corresponding coordinates contain the same information about [*X*]. However, the affine and projective features highlight different aspects of the EEG/MEG data frame *X* and one of them can be more natural than the other in a given setting.

### 4.1 Fundamental affine features

The fundamental affine features of a data frame *X* ∈ *F*_*n,k*_ are obtained by embedding the Grassmann manifold *G*_*n,k*_ in the vector space of real symmetric *n* × *n* matrices. This is done by mapping the *k*-plane [*X*] to the orthogonal projection matrix *P* (*X*) onto that *k*-plane:

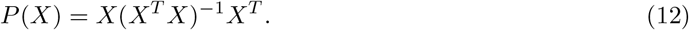

A more precise notation would be *P* ([*X*]) but we will use *P* (*X*) to emphasize that the projection matrix can directly be calculated from the data frame *X*. In order for the map *P* to be well-defined on *G*_*n,k*_, the matrix *P* (*X*) should not depend on which *k*-frame *X* is used to represent the *k*-plane [*X*]. So we need that *P* (*XA*) = *P* (*X*) for all real invertible *k* × *k* matrices *A*. That this is indeed the case follows directly from Eq. (12). Furthermore, the map *P* is indeed an embedding (i.e. is one-to-one), because any two different *k*-planes have different projection matrices. The image of *G*_*n,k*_ under this embedding is the set of *n*-dimensional projection matrices of rank *k*.

The Grassmann manifold *G*_*n,k*_ can hence be thought of as the (non-linear) subspace of the vector space of symmetric *n* × *n* matrices consisting of orthogonal projection matrices of rank *k*, or equivalently, with trace *k*:

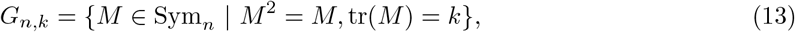

where Sym_*n*_ denotes the vector space of *n*-dimensional real symmetric matrices [24]. The dimension of Sym_*n*_ is *n*(*n* + 1)*/*2 and so the embedding can be used to associate with every EEG/MEG data matrix *X, n*(*n*+1)*/*2 invariant features, namely the upper-triangular entries of the matrix *P* (*X*). Since invariant features correspond to functions on *G*_*n,k*_, *any* invariant feature can be expressed in terms of these invariant features. We will refer to the upper-triangular entries of *P* (*X*) as the *fundamental affine features* of *X*.

What is the interpretation of the fundamental affine features in the context of EEG/MEG data? Let *X* be an *n* × *k* data frame with rows *x*^1^, …, *x*^*n*^. The vector *x*^*i*^ ∈ ℝ ^*k*^ is the spatial pattern (across the *k* brain locations) at time-point *i*. The (*i, j*)-th entry of *P* (*X*) is

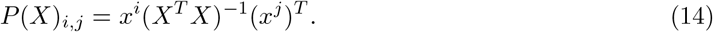

The fact that any invariant feature is a function of these entries is a classic result in statistics [34]. Eq. (14) shows that *P* (*X*)_*i,j*_ is the covariance between the spatial patterns at times *i* and *j* of the whitened data matrix. To make this more explicit, note that whitening of *X* does not change its column span: If *W* is a whitening transformation for *X* and *Y* = *XW* is the whitened data frame, then [*X*] = [*Y*], that is, *X* and *Y* correspond to the same point on the Grassmann manifold. We can therefore just as well work with *Y* so that Eq. (14) becomes

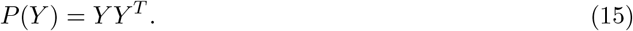

Write *y*^1^, …, *y*^*n*^ for the rows of *Y* and *y*_1_, …, *y*_*k*_ for its columns. So *y*^*i*^ ∈ ℝ ^*k*^ is the spatial pattern at time-point *i* of the whitened data matrix and *y*_*j*_ ∈ ℝ ^*n*^ is the observed signal at location *j*. The (*i, j*)-th entry

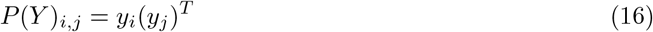

of *P* (*Y*) hence is the covariance between the patterns *y*^*i*^ and *y*^*j*^ at time-points *i* and *j*. We see that *P* (*Y*) is the temporal covariance matrix of the whitened EEG/MEG signals. In particular, its *i*-th diagonal entry 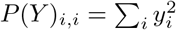 is the total power of the spatial pattern at time-point *i*. In Section 5.4 we will use the matrix *P* (*Y*) to construct invariant features of EEG/MEG data frames that relate to multivariate non-Gaussianity.

### 4.2 Fundamental projective features

The fundamental projective features of a data frame *X* ∈ *F*_*n,k*_ are obtained by embedding the Grassmann manifold *G*_*n,k*_ in the vector space ⋀ ^*k*^ ℝ ^*n*^ of alternating *k*-tensors on ℝ ^*n*^. The embedding map is known as the *Plücker embedding*. The image of a point [*X*] ∈ *G*_*n,k*_ is obtained by taking the wedge product of the columns of *X*:

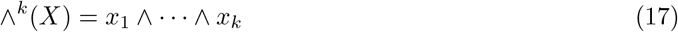

which is an alternating *k*-tensor on ℝ ^*n*^ (see Section 2.2). This image is only determined by [*X*] up to a scaling factor:

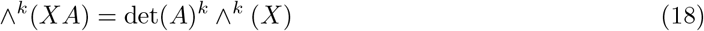

for all invertible *k* × *k* matrices *A*. To make the map well-defined, all scalar multiples of the tensor ∧^*k*^ (*X*) need to be considered as the same object. We need, in other words, to work with *lines* in ⋀ ^*k*^ ℝ ^*n*^, rather than with individual points. We thus need to work in the *projectivation* 𝒫 (⋀ ^*k*^ ℝ ^*n*^) of the vector space ⋀ ^*k*^ ℝ ^*n*^ (see Section 2.2). In the context of EEG/MEG data, we will refer to points on the embedded Grassmann manifold as *fundamental projective features*. Eq. (18) shows that the fundamental projective features are invariants of weight *k*.

As an example we consider the bivariate case (i.e. *k* = 2). Let *X* ∈ *F*_*n*,2_ be an EEG/MEG 2-frame. For clarity, we use a slightly different notation and denote the columns of *X* by *x* ∈ ℝ ^*n*^ and *y* ∈ ℝ ^*n*^. So *x* and *y* are observed EEG/MEG signals from two brain locations. The image of (*x, y*) under the Plücker embedding is

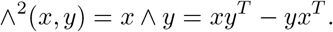

Its coordinates are *x*_*i*_*y*_*j*_ − *y*_*i*_*x*_*j*_, which are invariant features of weight 2. The product *x*_*i*_*y*_*j*_ can be thought of as the instantaneous linear interaction between the activity in the first location at time-point *i* and the activity in the second location at time-point *j*. Because *y*_*i*_*x*_*j*_ is obtained from *x*_*i*_*y*_*j*_ by interchanging the time-points *i* and *j, x*_*i*_*y*_*j*_ − *y*_*i*_*x*_*j*_ measures the lack of temporal reversibility in the interaction. Thus, a non-zero value of *x*_*i*_*y*_*j*_ − *y*_*i*_*x*_*j*_ reflects temporal irreversibility of the interaction at time-points *i* and *j*. It is closely related to the time-resolved connectivity measure proposed in [35].

As an illustration, consider the following simple model. Suppose *x* and *y* are harmonic oscillations of the same frequency *ω* rad*/*s and are sampled with *N* samples per second:

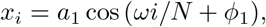

and

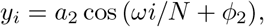

for *i* = 0, 1, 2, …, *n* and where *ϕ*_1_ and *ϕ*_2_ are phases. The Plücker embedding of (*x, y*) has the following coordinate at (*i, i* + *m*):

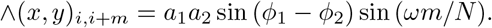

Note that the image does not depend on *i*, but only on the lag *m* = *j* − *i* and that the image vanishes if *ϕ*_1_ = *ϕ*_2_ (mod *π*), it vanishes. So if *x* and *y* are in-phase or in anti-phase, the image vanishes and hence it indeed measures the lack of temporal reversibility in the interaction at lag *m*,

We now consider the trivariate case (i.e. *k* = 3). Let *x, y, z* ∈ ℝ ^*n*^ be the observed signals from three different locations within the brain. The image of (*x, y, z*) under the Plücker embedding is the wedge product ∧^3^(*x, y, z*) = *x* ∧ *y* ∧ *z*. Its (*i, j, k*)-th coordinate is

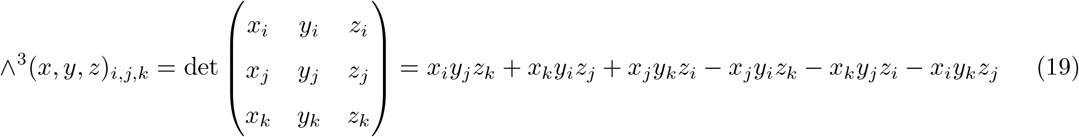

which is an invariant of weight 3. Observe that it involves *three* time-points *i, j*, and *k* and that it is constructed by summing triple instantaneous products of the signals at time-points *i, j*, and *k* over all six permutations of these three time-points, taking into account the signs of the permutations. For example, the product −*x*_*j*_*y*_*i*_*z*_*k*_ is obtained from *x*_*i*_*y*_*j*_*z*_*k*_ (the original ordering of the signals) by interchanging time-points *i* and *j*, which is a permutation with negative sign. This particular form is what makes it invariant under mixing. A non-zero value of ∧^3^(*x, y, z*)_*i,j,k*_ reflects the lack of symmetry of the instantaneous linear interaction at time-points *i, j*, and *k* under permutation of these time-points and hence is a measure of *temporal asymmetry*.

We now consider the general case. The fundamental projective invariants involving *k* signals *x*_1_, …, *x*_*k*_ have similar properties. The (*i*_1_, …, *i*_*k*_)-th entry of ∧^*k*^ (*x*_1_, …, *x*_*k*_) can be written as

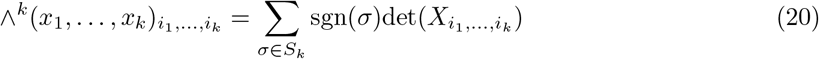

where 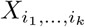 is the *k* × *k* submatrix of *X* obtained by selecting rows *i*_1_, …, *i*_*k*_ [36]. The entries of ∧^*k*^ (*x*_1_, …, *x*_*k*_) are (time-resolved) *k*-th order connectivity measures. A non-zero value of Eq. (20) reflects the lack of symmetry of the instantaneous linear interaction at time-points *i*_1_, …, *i*_*k*_ under permutation of these time-points and hence is a measure of *temporal asymmetry*. In particular, the fundamental projective invariants for *k* = 4 can be viewed as invariant versions of the recently proposed time-resolved edge connectivity measure in fMRI research [37, 38, 39, 40, 41].

In Section 5.2 we show how the fundamental projective invariants can be combined to build time-averaged higher-order connectivity measures that are invariant to linear and instantaneous mixing. The obtained measures can be used to study static higher-order brain interactions using EEG/MEG.

## 5 Construction of mixing-invariant EEG/MEG features

The fundamental projective features of an EEG/MEG data matrix containing two signals are instan-taneous bivariate connectivity measures and are hence suitable for time-resolved connectivity analyses. They are also relative invariants and hence can be used for hypothesis testing, but not for quantifying connectivity strength. In Section 5.1 we construct a bivariate connectivity measure of weight zero that can be applied to stationary signals. It is obtained by appropriate averaging and normalization of the fundamental projective features. In Section 5.2 we do this for more than two signals, thereby obtaining a higher-order connectivity measure of weight zero that can be applied to stationary signals.

### 5.1 Bivariate connectivity measures

Let *x, y* ∈ ℝ^*n*^ be observed EEG/MEG signals from two brain locations. The fundamental projective features of the pair (*x, y*) are the entries of the anti-symmetric *n* × *n* matrix

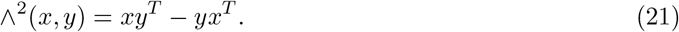

For a given lag *m >* 0, and typically much smaller than *n*, we construct a static connectivity measure *f*_*m*_ by averaging the entries ∧^2^(*x, y*)_*i,i*+*m*_ over time-points *i*:

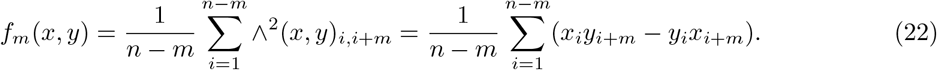

It measures the temporal irreversibility of the linear interaction between the signals *x* and *y* at lag *m* and can be used as a test-statistic for testing the null-hypothesis of temporal reversibility at lag *m*. It is, however, a relative invariant (it has weight 2) and therefore cannot be used to quantify the extent to which the signals are irreversible. But by dividing it by an invariant feature with the same weight, an absolute invariant connectivity measure is obtained as described below.

Let Σ be the 2 × 2 covariance matrix of the pair (*x, y*) and observe that 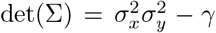 is an invariant of weight 4, so that

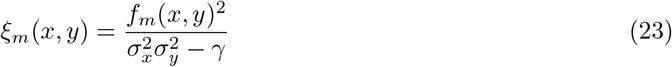

is an invariant connectivity measure of weight zero. Note that the measure in Eq. (23) is only well-defined if Σ is invertible. We can further combine the measures *ξ*_*m*_(*x, y*) by averaging over lags to obtain the following connectivity measure:

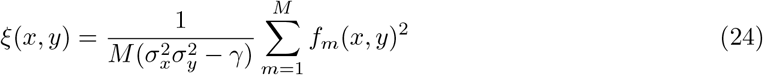

where *M* is the maximum lag to be taken into account. An appropriate value for *M* can be based on the characteristic time-scale of the covariance function *γ*_*m*_(*x, y*) of (*x, y*).

The above measure can be expressed in terms of the cross-correlation function of the pair (*x, y*) as follows. Recall that the cross-covariance function *γ*_*m*_(*x, y*) of the pair (*x, y*) is defined by

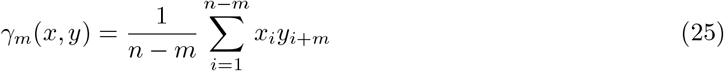

for *m* ≥ 0 and *γ*_*m*_(*x, y*) = *γ*_−*m*_(*y, x*) for *m <* 0. This implies

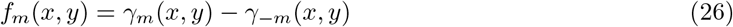

and hence

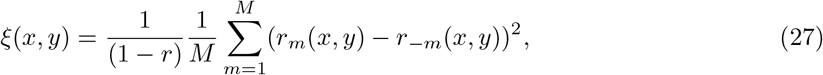

where *r*_*m*_(*x, y*) = *γ*_*m*_(*x, y*)/*σ*_*x*_σ_*y*_ is the cross-correlation function of (*x, y*) at lag *m* and *r* = *γ*_0_(*x, y*)/*σ*_*x*_*σ*_*y*_ is the instantaneous correlation between *x* and *y*. Eq. (27) shows that *ξ*(*x, y*) is a measure for the total asymmetry of the cross-correlation function of (*x, y*). In other words, *ξ*(*x, y*) is a measure for the *temporal irreversibility* of the linear interactions between *x* and *y*.

### 5.2 Third-order connectivity measures

In this section we consider the case of 3 signals. Let *x, y, z* ∈ ℝ^*n*^ be signals recorded from three locations within the brain. The fundamental projective features of the triplet (*x, y, z*) are the entries of the alternating 3-tensor ∧^3^(*x, y, z*). For given lags (*m*_1_, *m*_2_) with 0 ≤ *m*_1_, *m*_2_ ≤ *n* − *m*, where *m* = max{*m*_1_, *m*_2_}, we average the entries ∧^3^(*x, y, z*)_*i,i*+*m*, *i*+*m*_ over time-points *i* to obtain the following static third-order connectivity measure:

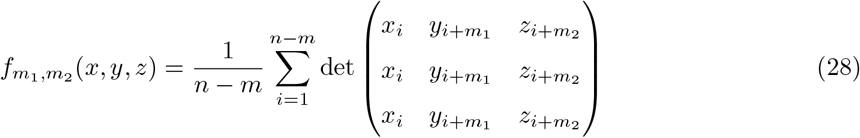

It measures the lack of symmetry of the linear interaction between the three signals at lags *m*_1_ and *m*_2_ under permutation of these lags and hence is a measure of *temporal asymmetry*. As in the bivariate case, this connectivity measure is a relative invariant (it has weight 3) but can be normalized to obtain an absolute invariant connectivity measure:

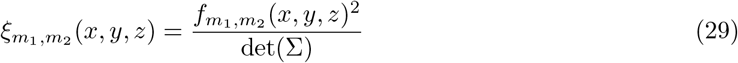

where Σ = *X*^*T*^ *X* is the covariance matrix of the *n* × 3 EEG/MEG data matrix *X* = [*x y z*], which is an invariant of weight six. Note that this measure is only defined if Σ is invertible. We can further combine the different lags by averaging to obtain the following connectivity measure:

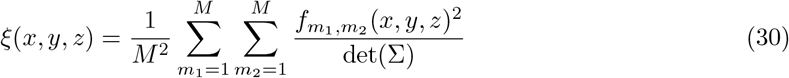

where *M* is the maximum lag to be taken into account. An appropriate value for *M* can be based on the characteristic time-scales of the covariance function 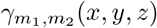

### 5.3 Multidimensional connectivity measures

In this section we describe how to construct invariant multidimensional connectivity measures between two data frames 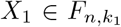 and 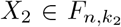 obtained from two brain regions, comprising *k*_1_ and *k*_2_ voxels. Invariance in this context pertains to mixing within each of the two regions (i.e. intra-regional mixing). Thus for a connectivity measure *f* (*X*_1_, *X*_2_) we require that

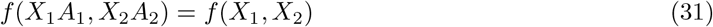

for all invertible matrices *A*_1_ and *A*_2_, which describe the mixing within the regions. To avoid introducing Cartesian products of Grassmann manifolds of different dimensions, we assume that the regions have the same number of voxels (i.e. *k*_1_ = *k*_2_ = *k*) but the formulas apply to the general situation as well.

Features that are invariant under intra-regional mixing characterize how pairs of *k*-planes (*V*_1_, *V*_2_) are embedded in ℝ*n*. Such features can be divided into two classes, namely those that only depend on the relative positions of the planes and those that also depend on the absolute positions of the planes. We focus on features that characterize relative positions. They correspond to symmetric (i.e. permutation-invariant) functions of the canonical angles between *V*_1_ and *V*_2_ (see Section 2.2) or, equivalently, to symmetric functions of the singular values of the spatial cross-covariance matrix 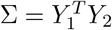 between the whitened frames *Y*_1_ and *Y*_2_. Note that such features are also invariant under reordering of the two regions (i.e. they are undirected).

Therefore, *any* function *f* that satisfies 𝔼[*f* (0)] = 0 is a functional connectivity measure between the multivariate activity in the two brain regions. This type of connectivity is referred to as *multidimensional* [17, 22, 23]. There exist infinitely many invariant multidimensional connectivity measures.

However, if we restrict to polynomial measures we can construct all of them by forming polynomial combinations of a finite set (of size *k*) of fundamental invariants. There exist several such fundamental sets, for instance, the elementary symmetric polynomials, the completely homogeneous symmetric polynomials, and the symmetric power sums in the singular values [42]. Each of these sets yields *k* algebraically independent multidimensional connectivity measures. From a geometric perspective, interesting choices are those that correspond to distance measures on the Grassmann manifold. An example is the geodesic distance

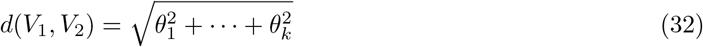

where *θ*_1_, …, *θ*_*k*_ are the canonical angles between *V*_1_ and *V*_2_. There are several more distance measures and each of them is a symmetric function of the canonical angles [43, 24].

### 5.4 Features based on spatial activation patterns

Let *X* be an *n* × *k* EEG/MEG data frame and let *Y* be a whitened version of it. The matrix *Y* can be taken to be the *n* × *k* matrix of left singular vectors of *X*. The rows of *Y* are the spatial patterns of the whitened EEG/MEG signals at the *n* different time-points and they are denoted by *y*^1^, …, *y*^*n*^. Recall from Section 4.1 that the fundamental affine features of *X* are the entries of the temporal covariance matrix *P* (*Y*) = *Y Y* ^*T*^. Thus, the (*i, j*)-th entry *P*_*i,j*_ = ⟨ *y*^*i*^, *y*^*j*^ ⟩ is the covariance between the spatial patterns *y*^*i*^ and *y*^*j*^ at time-points *i* and *j*. Below we describe several ways to combine these covariances to construct invariant features for characterizing the spatiotemporal dynamics within an EEG/MEG data frame *X*.

If we divide the (*i, j*)-th entry of *P* (*Y*) by the product of the norms ||*y*^*i*^|| and ||*y*^*j*^|| of *y*^*i*^ and *y*^*j*^, we obtain the *temporal correlation matrix C*(*Y*) of *Y*. Its (*i, j*)-th entry

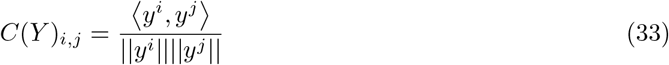

is the correlation between the spatial patterns *y*^*i*^ and *y*^*j*^ at time-points *i* and *j*. The temporal correlation matrix *C*(*Y*) characterizes the dynamics of the EEG/MEG signals for each pair of time-points and hence is time-resolved and suitable for dynamic analyses of non-stationary brain activity. If the EEG/MEG signals are assumed to be stationary, the entries of the temporal correlation matrix can be averaged over the super-diagonals, to obtain correlation coefficients that only depend on the time-lag *i*−*j* between *y*^*i*^ and *y*^*j*^. Another way of reducing the information contained in the temporal correlation matrix is to calculate a matrix norm (e.g. the Frobenius norm) its determinant, or by combining its entries in any other way (e.g. by averaging over columns).

We can also use the fundamental affine features to construct dissimilarity measures between the spatial patterns *y*^*i*^ and *y*^*j*^ at two different time-points *i* and *j*. A general way of constructing such measures is by defining a distance *D*(*Y*)_*i,j*_ between the vectors *y*^*i*^ and *y*^*j*^. Any distance that is a function of the covariances ⟨ *y*^*i*^, *y*^*j*^ ⟩ is invariant under mixing. For example, the Euclidean distance *D*(*Y*)_*i,j*_ = ||*y*^*i*^−*y*^*j*^|| between *y*^*i*^ and *y*^*j*^ can be expressed in terms of covariances as

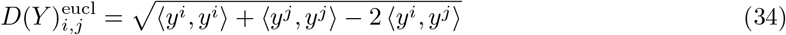

and hence is invariant. Another example is the cosine distance between *y*^*i*^ and *y*^*j*^, which is defined by

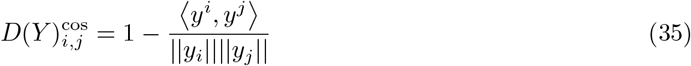

and hence is invariant under mixing. Note that the cosine distance between *y*^*i*^ and *y*^*j*^ equals one minus the temporal correlation between *y*_*i*_ and *y*_*j*_. As with the temporal correlation matrix *C*(*Y*), the information contained in dissimilarity matrices can be reduced by averaging over super-diagonals, by calculating the matrix norm, or by combining its entries in any other way.

An interesting invariant feature is obtained by averaging the diagonal entries of the third tensor power *P* (*Y*) ⊗ *P* (*Y*) ⊗ *P* (*Y*) over time-points (i.e. taking its trace and dividing by *n*). This produces the following feature:

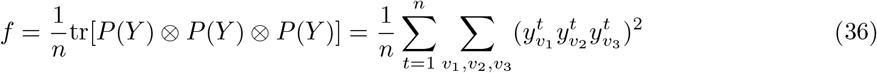

where *v*_1_, *v*_2_, *v*_3_ run over all *k* brain regions. This is the multivariate skewness measure proposed in [18] and can be used to quantify deviations from multivariate normality. A related measure is the multivariate kurtosis [18] and it can be expressed in a similar way as a function of the diagonal entries of the fourth tensor power of *P* (*Y*). Whereas the multivariate skewness is sensitive to the third-order statistical moments of the signals (coskewness), the multivariate kurtosis is sensitive to the fourth-order statistical moments of the signals (cokurtosis). Related measures for multivariate non-Gaussianity can be found in [44].

## 6 Application to EEG/MEG data

Our methods could be fruitful for the EEG/MEG connectivity field if our metrics provide complementary information to existing measures. To illustrate their potential, we apply the temporal irreversibility metric (Eq. (27)) to clinical EEG data. Pathological conditions are especially suitable for testing such propositions, as they allow for testing sensitivity to disease states.

A large body of literature has shown that EEG monitoring allows early and reliable prediction of good or poor recovery of comatose patients after cardiac arrest [45, 46, 47, 48]. We include EEG data from hundred patients, as described in our previous work [49]. All patients underwent continuous EEG monitoring (standard 10-20 system) as part of routine clinical care. Details of data acquisition and inclusion criteria are also reported in [49]. In brief, we included fifty patients with a poor neurological outcome and fifty patients with a good neurological outcome, defined by the cerebral performance category (CPC) score, assessed six months after cardiac arrest. The CPC score quantifies the neurological outcome on a 5-point scale. A CPC=1 (no cerebral damage) or CPC=2 (moderate disability) is generally considered a favorable outcome, while the scores CPC=3 (severe disability), CPC=4 (minimal conscious state or non-responsive sleep-wake syndrome) or CPC=5 (death) reflect a poor neurological outcome.

We selected five minutes of EEG data around 48 hours after cardiac arrest in patients who had developed a discontinuous or continuous EEG at this interval. Artefacts were excluded using an automated custom computer algorithm that rejected flat channels, sensor noise, and muscle artefacts [50]. In addition, we used independent component analysis (ICA) to identify electrocardiogram artefacts [51]. Every ICA component with a strong correlation with the electrocardiogram signal (defined as a Pearson correlation greater than 0.6) was rejected. This resulted in 0-2 rejected ICA components for every subject. Artefact rejection was followed by source reconstruction of the EEG data using *exact low-resolution brain electromagnetic tomography* (eLORETA) [52] as implemented in Fieldtrip [53]. A template T1-weighted image was used to compute a Boundary Element Method (BEM) head model for every subject [54]. Source reconstructed data were parcellated using the automated anatomical atlas (AAL) [55], and time-series within a parcel were averaged to obtain a single time-series per parcel. Only cortical parcels were included for further analysis [56], resulting in 78 virtual electrodes. Source reconstructed data were band-pass filtered using a finite impulse response (FIR) filter into the canonical frequency bands: delta (1-4 Hz), theta (4-8 Hz), and alpha (8-13 Hz). For every frequency band, we computed phase synchrony using an existing measure that is invariant to signal leakage, i.e. the phase lag index (PLI) [14]. In addition, we computed the temporal irreversibility metric (Eq. (27), with a maximal lag of a second) to the band-pass filtered signals recorded from the virtual electrodes. Statistical testing between groups was performed using *t*-tests with correction for multiple comparisons using the false discovery rate [57].

Figure 8-A and Figure 8-B show the group-averaged temporal irreversibility matrix and corresponding connectivity maps for patients with good and poor outcomes for the delta band respectively. In the poor outcome group, there is overall stronger temporal irreversibility in the delta band across many brain areas. This is also confirmed by formal statistical testing as demonstrated in Figure 9C, with group differences in the strength of temporal irreversibility especially between left and right temporal and motor areas. Note that these statistically significant group differences were more pronounced than for the phase lag index (PLI) for the same data (Figure 9D). In fact, we observe that regions for which the PLI showed significant effects are a subset of the regions that are significantly different for temporal irreversibility. Hence, this suggests that our invariant temporal irreversibility is more sensitive to detecting disease-induced effects. There were no significantly different results for temporal irreversibility for other frequency bands.

**Figure 8.**
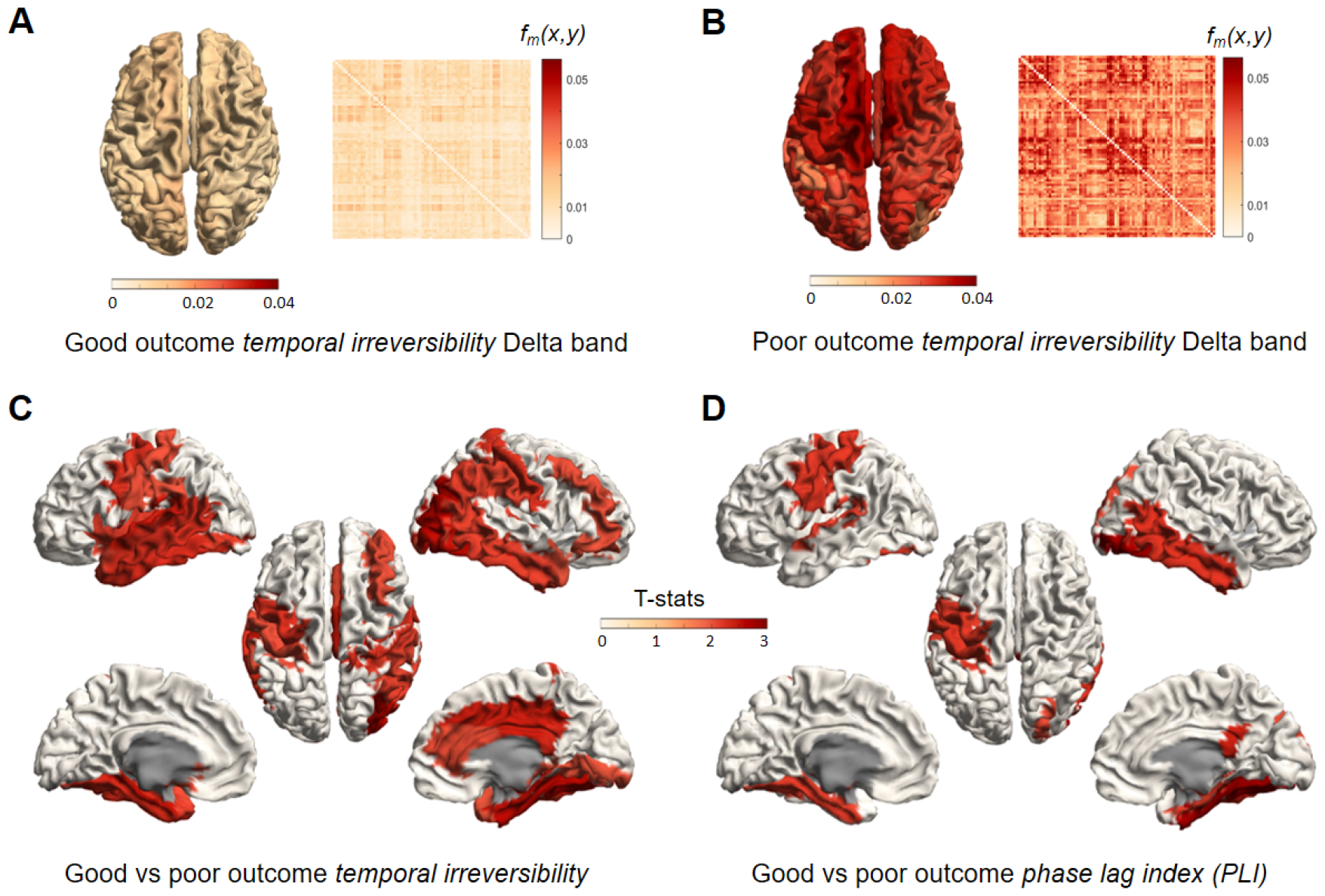
Temporal irreversibility (Eq. (27)) applied to EEG data of comatose survivors of cardiac arrest. Panels A and B show the average temporal irreversibility matrix across patients in each group. The column average of these matrices is plotted next to them as a brain plot. There is much stronger temporal irreversibility in patients with a poor outcome. Panels C and D show the outcome of statistical testing for the temporal irreversibility and phase lag index, respectively. Colours represent *t*-values for regions that show significant differences between groups (*p <* 0.05 FDR corrected).sss

## 7 Conclusions and discussion

In this study we have described two sets of fundamental features of EEG/MEG data matrices that are invariant under linear and instantaneous mixing and from which all other invariant features can be constructed by combination, namely the affine and projective features. These two sets of features can hence be viewed as building blocks of invariant features. We have illustrated their usefulness by constructing several new invariant features of EEG/MEG data matrices, including bivariate, higher-order, and multidimensional connectivity measures. They were derived by exploiting the correspondence between invariant features and functions on the Grassmann manifold and subsequently using the topological and geometric properties of the Grassmann manifold. The main advantage of invariant features over general (non-invariant) features, is that they allow for a functional interpretation because they are not subject to signal leakage. We expect that our results will be useful in general analyses of EEG/MEG data and in classification and prediction problems using machine learning in particular. For instance, the fundamental features can be used as features in classification or prediction tasks.

In our study, invariance of the derived features was guaranteed by working directly with planes (i.e. points on the Grassmann manifold) instead of with frames. This approach can be generalized to other analyses of EEG/MEG data and hence allows designing signal processing pipelines, the results of which are invariant by construction. For example, averaging of EEG/MEG data matrices (over subjects, conditions, etc.) can be done on the Grassmann manifold [58] and ensures that the average matrices have full column-rank. This approach is in the same spirit as using the manifold of symmetric positive-definite (SPD) matrices for averaging covariance matrices, which preserves positive-definiteness [59]. The manifold of SPD matrices has been proposed as a general framework for carrying out functional connectivity analyses of fMRI data [60, 61, 59]. Other methods that are available for Grassmann-valued data are linear regression, clustering, and kernel methods [62, 58, 63].

In discussing multidimensional connectivity measures, we assumed both regions to have the same number of voxels. This assumption, however, is not essential and the results are applicable to regions with different numbers of voxels. We made this assumption to keep the mathematical formalism to a minimum: If the regions have the same number of voxels, the corresponding data frames correspond to points on the Grassmann manifold, whereas if they have different numbers of voxels, we would have to discuss products of Grassmann manifolds of different dimensions, which requires additional concepts from algebraic geometry. A general theory for products of Grassmann manifolds is worked out in [43]. Since the theory applies to products of arbitrary numbers of Grassmann manifolds, it can be used to generalize multidimensional connectivity measures to more than two brain regions of possibly different numbers of voxels. This would lead to connectivity measures that are both multidimensional and higher-order.

In discussing multidimensional connectivity measures, the data frames were thought to correspond to different brain regions. However, they can also thought to correspond to different trials, subjects, groups, or conditions. In these applications, it is evident that there is no mixing between the frames, since they are not obtained from simultaneous recordings of single subjects. If the frames are obtained from simultaneous recordings of single subjects, there might be mixing between the frames, depending on the exact brain regions. If the reconstructed source signals are obtained from a linear inverse operator, such as the minimum norm solution [10] or a linearly constrained minimum variance beamformer [26, 27], the mixing of sources is described by the resolution operator, which is the concatenation of the forward and inverse operators [28, 29]. Inspection of the resolution operator then allows assessing the severity of mixing between the regions [23]. Although this does not solve the mixing problem, analyses can at least be restricted to regions with minimal mixing.

We have applied one of the Grassmann-derived invariant measures, the temporal irreversibility metric, to EEG data of comatose survivors of cardiac arrest. We demonstrated that this metric is more sensitive to identifying affected brain regions between patients with a good and poor outcome than a conventional invariant phase synchrony measure, the phase lag index [64]. Remarkably, regions identified with our metric include the subset of regions that were identified with the phase lag index. While the usage of asymmetry in connectivity may not only be justified by its insensitivity to signal leakage, it may also be more sensitive to long-distance communication between neuronal populations. Genuine connectivity between populations is associated with a conduction delay between populations and a presumed preferred direction of connectivity. The temporal irreversibility metric may be more sensitive to this characteristic than conventional phase synchrony, which does not distinguish consistent intervals of phase lag versus phase lead. We note that the temporal irreversibility metric is strongly related to the irreversibility metric introduced by the so-called “inside out framework” as introduced by Deco and coworkers [65]. In line with previous work [66], we have demonstrated that at 48 hours after cardiac arrest, there were only significant differences in temporal irreversibility in the delta band between patients with a good and poor neurological outcome. Presumably, an increased functional connectivity of delta oscillations at this time point after arrest is associated with poor neurological recovery. Further clinical research in larger clinical cohorts is warranted to study the additional value of irreversibility in the delta band along existing prognostic EEG features in comatose patients after cardiac arrest.

## Acknowledgements

R.H. was funded by NWO-Wiskundeclusters grant nr. 613.009.105.

## Author contributions

RH and TR designed the study, RH and PT wrote the manuscript, and MP and PT carried out the data analysis.

## Data Availability Statement

All EEG data and Matlab code are available on Zenodo at https://zenodo.org/doi/10.5281/zenodo.10700040.

